# Divergent Entry, Convergent Trafficking in Defensin-Mediated Adenovirus Infection

**DOI:** 10.64898/2026.08.13.744638

**Authors:** Cheng Zhao, Jason G. Smith

## Abstract

Human α-defensins can paradoxically inhibit or enhance human adenovirus (HAdV) infection. Defensin-resistant HAdVs exploit α-defensins as molecular bridges to bypass canonical receptors, but whether this mode of entry redirects intracellular trafficking or contributes to defensin resistance remains unknown. Here we systematically compared intracellular trafficking after canonical receptor-mediated versus defensin-mediated entry using a defensin-resistant chimeric virus based on HAdV-C5 together with the human α-defensins, HD5 and HNP1. Pharmacological perturbations, capsid mutants, and imaging revealed that both entry routes rapidly converge on common checkpoints. Defensin-mediated infection showed the same sensitivity to endosomal acidification blockade and comparable co-localization with early endosomes as canonical entry. Notably, the membrane-lytic activity of HD5 did not substitute for viral protein VI, indicating that endosomal escape still requires intact protein VI function. Downstream transport also overlapped, as defensin-mediated infection, like canonical entry, depended on dynein-driven microtubule transport and passage through the microtubule-organizing center (MTOC) for nuclear entry. Co-infection experiments further showed that defensin-sensitive and defensin-resistant viruses retain their intrinsic phenotypes within the same cell, indicating that defensin effects are virion-autonomous and determined at or before cell entry rather than by downstream trafficking. These findings support a model in which resistance to α-defensin neutralization is governed by capsid features that control defensin binding and its effects on uncoating, not by access to an alternative intracellular pathway.

**Author Summary:** Human adenoviruses can be neutralized by α-defensins, antimicrobial peptides that bind the capsid and block uncoating. Paradoxically, some defensin-resistant adenoviruses infect cells more efficiently in the presence of defensins, with the peptides acting as molecular bridges that allow the virus to attach independently of its normal receptors. We asked whether this alternative attachment route changes how adenoviruses traffic through the cell, which could contribute to defensin resistance.

Using labeled viruses, inhibitors, capsid mutants, and receptor knockout cells, we compared trafficking after canonical versus defensin-mediated entry. Both routes converged at the earliest steps of infection, with identical endosomal acidification requirements, similar early endosome colocalization, and the same dependence on microtubule transport to the nucleus. Co-infection experiments demonstrated that sensitive and resistant viruses retained distinct phenotypes in the same cell, indicating susceptibility is set at or before entry.

These results demonstrate that defensin effects are decided at the plasma membrane, where capsid features, not alternative entry routes, drive resistance. Adenovirus entry is flexible at the cell surface, where the virus can attach through multiple mechanisms, but narrows to a single, constrained trafficking route inside the cell. Our findings inform how defensin-resistant vectors may behave in defensin-rich tissues during gene therapy or virotherapy.

## Introduction

Human adenoviruses (HAdVs) are non-enveloped, double-stranded DNA viruses of the *Adenoviridae* family that cause a wide spectrum of clinical diseases, including respiratory tract infections, gastroenteritis, conjunctivitis, and severe disseminated infections in immunocompromised individuals [1–3]. The adenovirus infection cycle begins with viral attachment to specific cellular receptors [e.g., CAR (coxsackievirus and adenovirus receptor) or CD46] via the viral fiber protein [4]. Partial uncoating at the cell surface followed by internalization through clathrin-mediated endocytosis is facilitated by interactions between the penton base protein and integrin co-receptors [5,6]. After entry into the host cell, the endosomal environment induces further disassembly of the viral capsid and release of the membrane-lytic protein VI, enabling the subviral particle to escape into the cytoplasm [7]. The virus is then transported along microtubules to the perinuclear microtubule-organizing center (MTOC), dislodged from the MTOC, and re-localized to the nuclear pore complex (NPC) [6]. Further uncoating occurs at the NPC, and the viral genome is delivered into the nucleus. There, transcription and replication proceed using both host and viral factors, ultimately leading to the production of progeny virions and cell lysis [8].

Human α-defensins, a class of cationic antimicrobial peptides comprising myeloid human neutrophil peptides 1 to 4 (HNP1 to HNP4) and enteric α-defensins HD5 and HD6, have emerged as key modulators of adenovirus infection [9–15]. Widely recognized for their broad-spectrum activity against bacteria via membrane disruption and other cytolytic mechanisms, α-defensins have also been shown to possess potent antiviral activity against both enveloped and non-enveloped viruses including HAdVs [16–22]. α-defensins bind directly to the adenovirus capsid, and neutralization determinants lie both in the hexon proteins that constitute the bulk of the icosahedral facets and in the vertex region composed of fiber and penton base [11,13,15,23–25]. This binding forms a molecular clamp that stabilizes the capsid and prevents the conformational changes necessary for uncoating and protein VI release, thereby blocking endosomal escape and subsequent genome delivery to the nucleus. Capsid modifications that alter these determinants, either naturally occurring or engineered, allow some serotypes or mutants of HAdVs to resist or evade the α-defensin uncoating block.

Paradoxically, the infectivity of some defensin-resistant HAdVs is enhanced by α-defensins [9,11,13]. This enhancement occurs through a mechanism in which α-defensins act as molecular adapters, bridging viral particles to host cells independently of canonical receptors [23]. In so doing, α-defensins facilitate attachment to otherwise non-susceptible cells and broaden adenovirus tropism; however, the mechanisms that govern intracellular virus transport following defensin-mediated entry and the potential for differential trafficking to contribute to defensin resistance remain poorly understood. In this study, we systematically dissect the intracellular trafficking of HAdV following canonical, receptor-mediated and non-canonical, defensin-mediated entry using a combination of pharmacological inhibitors, genetically modified cell lines, and capsid-mutant viruses. Our results demonstrate that while α-defensins facilitate HAdV uptake, they do not allow the virus to bypass canonical trafficking steps. Instead, defensin-mediated infection proceeds through a pathway that remains dependent on endosomal trafficking and microtubule transport. We also find no evidence that the inherent membrane lytic activity of α-defensins contributes to HAdV endosomal escape. Co-infection experiments further indicate that defensin-sensitive and defensin-resistant viruses retain their intrinsic behaviors within the same cell, consistent with virion-autonomous effects established at entry. Thus, defensin resistance does not arise from differential intracellular routing but from capsid features that dictate how α-defensins influence infection.

## Results

### Canonical and defensin-mediated HAdV entry utilize a common endosomal route

In the canonical entry pathway, HAdV is endocytosed after receptor binding, which is facilitated by integrin-mediated signaling [26]. Penton base/integrin interactions are also required for productive α-defensin-mediated infection, suggesting a common entry mechanism [23]. To compare receptor-and α-defensin-mediated entry, we used the previously published C5/D64-HVR1 chimera virus [23], which was engineered by replacing the hexon hypervariable region 1 (HVR1) of C5 with that of the naturally defensin-resistant D64, thereby conferring α-defensin resistance. We then established three routes of infection: 1) entry of C5/D64-HVR1 into WT A549 cells in the absence of HD5 is solely CAR-mediated, 2) entry of C5/D64-HVR1 into CAR knockout (KO) A549 cells in the presence of HD5 is solely α-defensin-mediated, and 3) entry of C5/D64-HVR1 into WT A549 cells in the presence of HD5 may use both routes.

To assess endocytosis, we first examined the effects of a chemical inhibitor on infection. Although there is evidence to suggest that HAdVs exit early endosomes and are not exposed to the acidified environment of late endosomes, HAdV entry paradoxically remains sensitive to a variety of reagents that disrupt endosomal acidification [6,27]. These agents include ionophores, inhibitors of the vacuolar H^+^-ATPase, and weak bases such as ammonium chloride (NH₄Cl). We therefore treated cells with 50 mM NH₄Cl and assessed C5/D64-HVR1 infection. We also included EV-A71 as a positive control virus with known sensitivity to inhibition by NH₄Cl [28]. We found that NH₄Cl treatment prior to infection significantly inhibited HAdV infection by both routes (Fig 1A).

**Fig 1.**
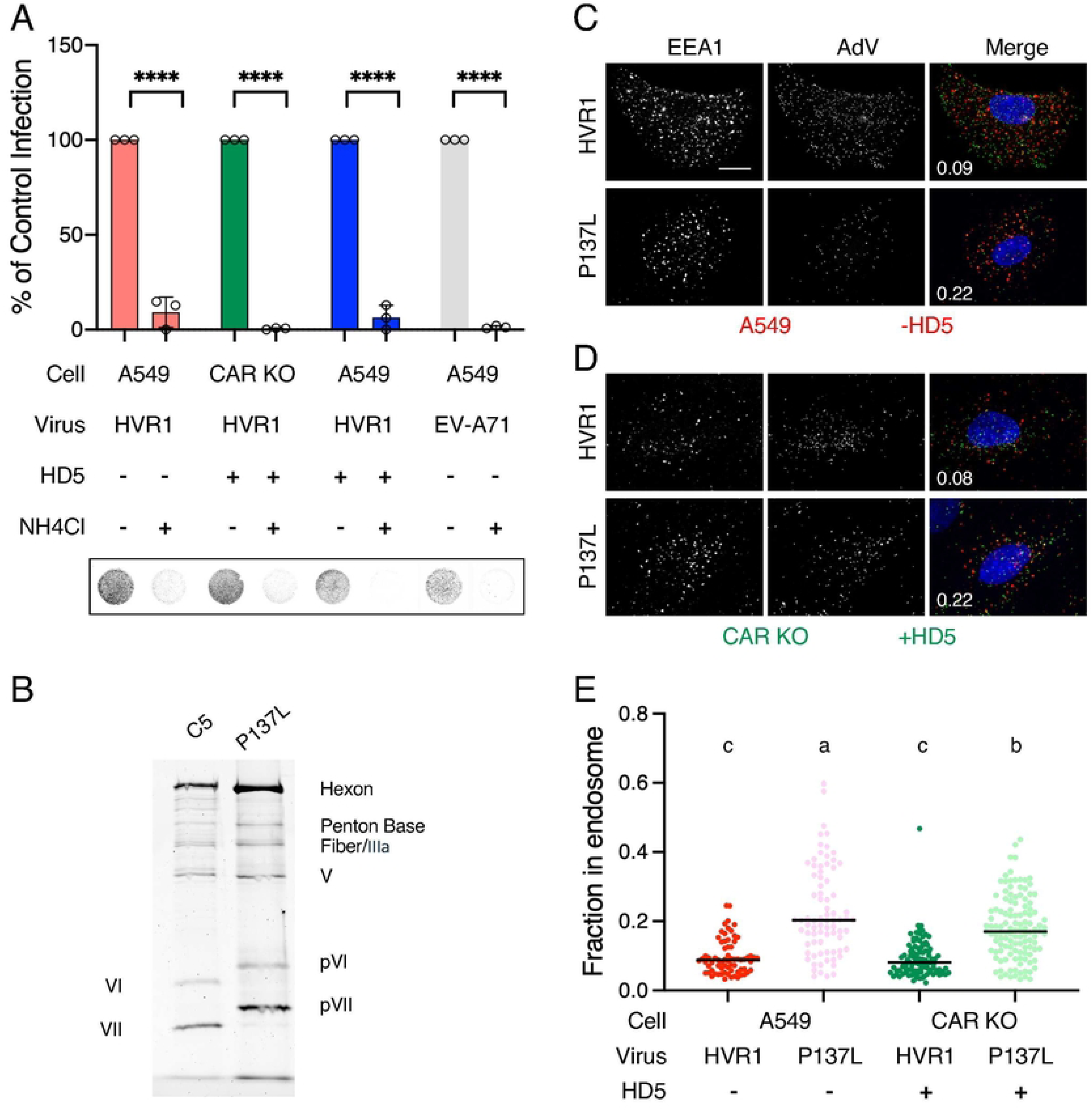
HD5-mediated entry of HAdV is via an ammonium chloride-sensitive endosomal route. (A) WT and CAR KO A549 cells were infected with either the C5/D64-HVR1 chimera virus or EV-A71 control virus with or without 5 µM HD5. Where indicated, cells were pretreated with NH₄Cl for 1 h prior to infection, and the inhibitor remained present for an additional 2 h p.i. Each point is an independent replicate (n = 3), and bars are the means ± SD of the percent infectivity compared to untreated control cells. Data were analyzed by ordinary two-way ANOVA with Šídák’s multiple comparisons test with a single pooled variance comparing infection with and without NH_4_Cl for each virus. ****, *P* < 0.0001. Representative wells of a 96-well plate are shown below, where grayscale intensity correlates with eGFP expression (C5/D64-HVR1) or immunofluorescence (EV-A71), indicative of successful infection. (B) Purified C5/D64-HVR1 and the protease P137L mutant propagated at the non-permissive temperature (39.5°C) were analyzed by SDS-PAGE and AzureRed staining. Bands corresponding to hexon; penton base; fiber; proteins IIIa, V, VI, and VII; and the precursors of proteins VI (pVI) and VII (pVII) are indicated. Representative images of (C) WT or (D) CAR KO A549 cells infected with Alex Fluor 488 (AF488)-labeled C5/D64-HVR1 or the protease P137L mutant virus in the (C) absence or (D) presence of 5 µM HD5. Images are maximum intensity Z-projections from 30 min p.i. Merged images include virus (AF488, green), a marker of early endosomes (EEA1, red), and the nucleus (DAPI, blue). Scale bars indicate 10 μm. The fraction of virus colocalized with EEA1 is indicated for each image. (E) Colocalization of virus with EEA1 was assessed for the indicated combinations of viruses, cells, and HD5 30 min p.i. by confocal microscopy and automated image analysis. Each point is an individual cell (n>70) from one of three biological replicates that were pooled and analyzed in aggregate. Horizontal bars indicate means. Data were analyzed by ordinary one-way ANOVA with Tukey’s multiple comparisons test with a single pooled variance comparing the mean of each column with every other column. Different letters indicate significant differences between groups (*P* < 0.05). Groups that share a letter are not statistically significantly different from one another.

To directly observe viral localization within cellular endosomes, we performed immunofluorescence staining. Because HAdV trafficking in endosomes is transient, we created a virus defective for endosomal escape by engineering a mutation (P137L) in the protease of C5/D64-HVR1. This mutation was first discovered in the *ts1* mutant of HAdV-C2 [29]. It blocks incorporation of the viral protease into assembling virions, thereby preventing maturational cleavage of the capsid proteins [30]. This is indicated by the presence of the precursor forms of protein VI and VII (pVI and pVII, respectively) in purified protease P137L mutant virions (Fig 1B). Accordingly, protease P137L mutant virions fail to properly uncoat in the endosomal system and release mature protein VI, which is required for endosomal escape [31–33]. We conjugated Alexa Fluor 488 (AF488) to the protease P137L mutant and parent virus, infected cells with the labeled viruses, and stained for the early endosomal marker EEA1 at 30 min post-infection (p.i.). Compared to the parent C5/D64-HVR1 chimera, colocalization of the protease P137L mutant with EEA1 was ∼2.5-fold higher. This was true for both the canonical entry route (Fig 1C and 1E) and the α-defensin-dependent pathway (Fig 1D and 1E). Taken together, these results suggest that α-defensin-mediated entry follows an endosomal route comparable to that of canonical entry.

### HD5 membrane lytic activity does not facilitate HAdV endosomal escape

Since α-defensins are membrane lytic molecules, we considered the possibility that HD5 co-internalized with HAdV could facilitate endosomal escape by functionally replacing or complementing the activity of protein VI. Such a mechanism might contribute to α-defensin-mediated enhanced infection at a second stage in entry beyond receptor binding. To test this, we created a second mutant virus by engineering a mutation (L40Q) in protein VI of the C5/D64-HVR1 chimera (Fig 2A). Unlike the protease P137L mutant virus described above, viruses bearing VI L40Q uncoat normally; however, the activity of the N-terminal amphipathic helix of protein VI is abrogated by the mutation, and these viruses are defective in endosomal escape [7].

**Fig 2.**
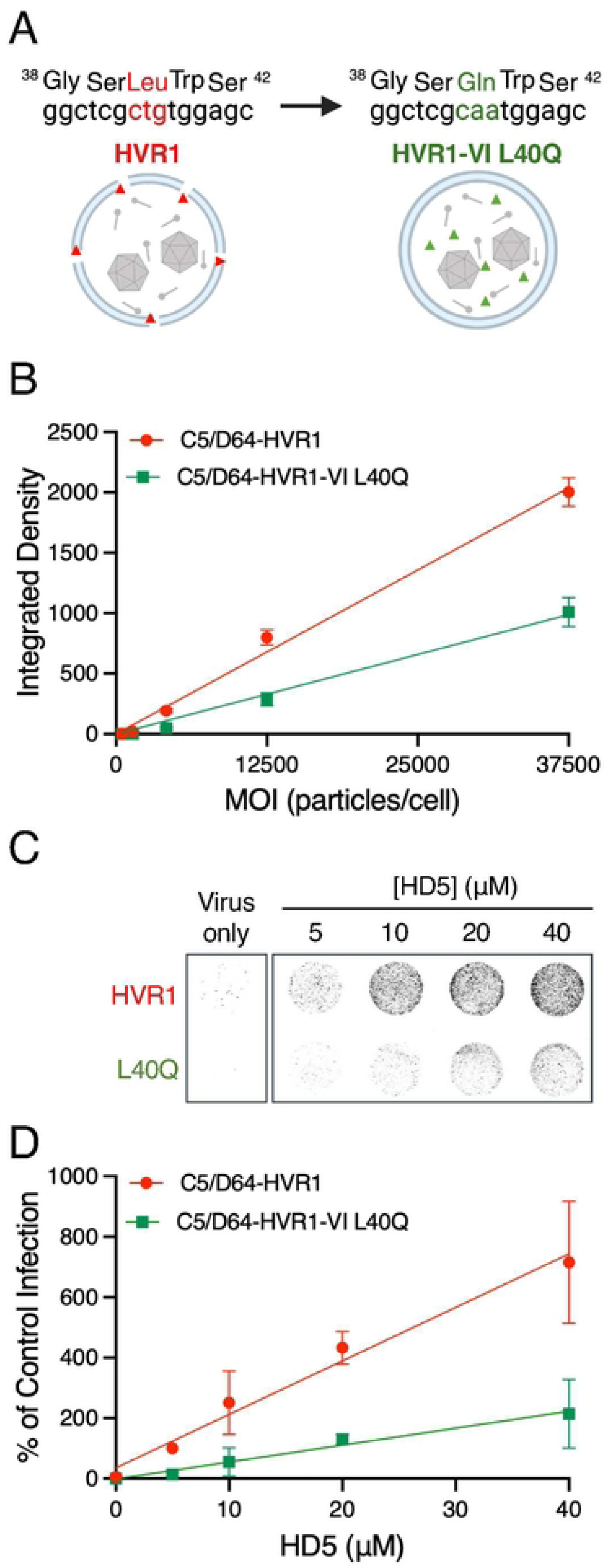
HD5 does not rescue a HAdV mutant defective in endosomal escape. (A) Schematic of the VI L40Q mutation that renders C5/D64-HVR1 defective in endosome escape. Figure created with BioRender.com. (B) A549 cells were infected with serial dilutions of the parent C5/D64-HVR1 or VI L40Q mutants at equivalent MOIs by particle number. Data are the means ± SD of the integrated densities of the background-subtracted eGFP signal of the cell monolayers from 3 independent replicates. (C) Representative images and (D) quantification of infection of CAR KO A549 cells with the indicated viruses in the presence or absence of HD5. Images in C were obtained at a resolution of 50 μm, and grayscale intensity correlates with eGFP expression. In D, data are normalized to C5/D64-HVR1 infection in the presence of 5 μM defensin (control infection) and are the means ± SD of 3 independent replicates.

To verify the mutant virus phenotype, we first infected WT A549 cells with both the VI L40Q mutant and parent C5/D64-HVR1 chimeras at comparable viral particle to cell ratios (particle MOI). As expected, the infectivity of the VI L40Q on a per particle basis was reduced compared to the parent virus (Fig 2B). We then infected CAR KO A549 cells with both viruses at the same particle MOI but in the presence of increasing concentrations of HD5. In the absence of HD5, C5/D64-HVR1 infection was barely detectable, and this residual infectivity was further reduced for the VI L40Q mutant. However, infectivity of both viruses was restored in the presence of HD5 (Fig 2C and 2D). For the parent C5/D64-HVR1 chimera, we observed a linear relationship between infection and HD5 concentration. We hypothesized that if HD5 promoted endosomal escape in addition to cell binding, the infectivity dose-response of the VI L40Q virus would curve upwards with increasing HD5 concentration; however, the VI L40Q mutant had both lower infectivity at each HD5 concentration than the parent virus and its infectivity exhibited linear dependence on defensin concentration (Fig 2D). Moreover, the ratio of the slopes of the best-fit infectivity lines of the two viruses as a function of HD5 concentration (Fig 2D, C5/D64-HVR1, slope = 0.054; VI L40Q mutant, slope = 0.026; ratio = 2.08) mirrors that of the best-fit infectivity lines of the two viruses as a function of MOI (Fig 2B, C5/D64-HVR1, slope = 1.80; VI L40Q mutant, slope = 0.92; ratio = 1.97). This suggests that the main effect of HD5 on promoting α-defensin-resistant virus entry is to mediate cell binding rather than also exerting membrane lytic activity and promoting viral endosomal escape.

### Defensin-mediated HAdV infection depends on microtubule transport

Following classical receptor-mediated HAdV entry and endosomal escape, viruses are transported through the cytoplasm to the nucleus via microtubules [34]. A balance between dynein-and kinesin-mediated transport is required for the virus to approach the nucleus. To investigate whether α-defensin-mediated viral infection also utilizes microtubule transport for cytoplasmic trafficking, we used the dynein inhibitor Ciliobrevin D [35]. We first determined the pre-treatment time and concentration of Ciliobrevin D required to block C5/D64-HVR1 infection of WT A549 cells. We found that 67 μM Ciliobrevin D treatment of A549 cells for 4 h inhibited infection by 75% (Fig 3A-B), and this concentration was chosen for subsequent experiments. We then determined the effect of Ciliobrevin D on canonical and HD5-mediated infection. We again included EV-A71 infection of WT A549 cells as a control because EV-A71 replicates in the cytoplasm and does not require microtubule-dependent nuclear transport [36]. Ciliobrevin D treatment significantly inhibited C5/D64-HVR1 infection by both routes but had no effect on EV-A71 infection (Fig 3C). The lack of inhibition of EV-A71 suggests that Ciliobrevin D treatment is not generally cytotoxic and that the effect on C5/D64-HVR1 infection is due to a block in viral trafficking rather than a disruption of the endosomal system, since EV-A71 infection requires acidified endosomes (Fig 1A). Thus, like canonical infection, α-defensin-promoted infection depends on microtubule transport.

**Fig 3.**
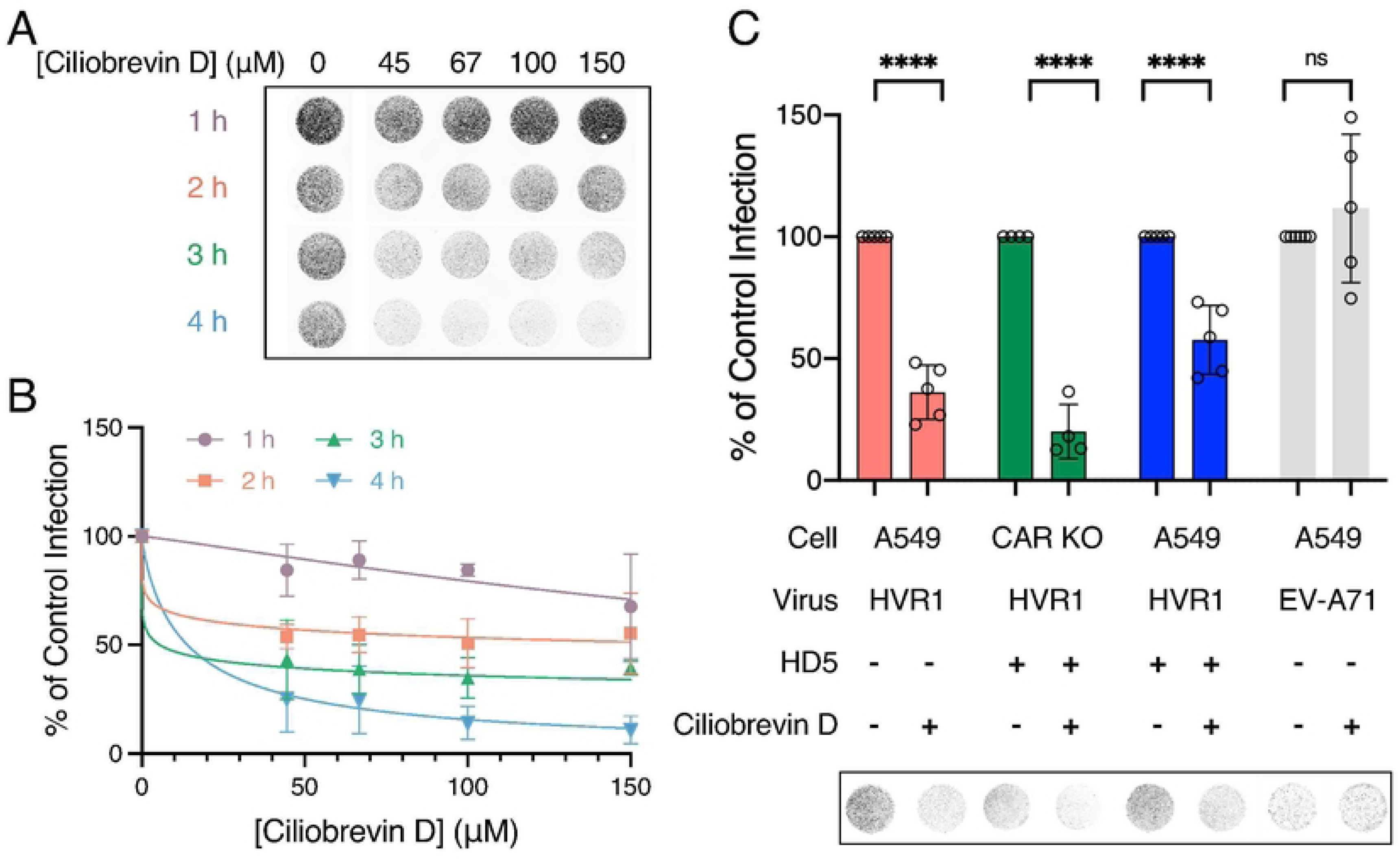
HD5-mediated HAdV entry is dynein-dependent. (A) Representative images and B) quantification of infection of A549 cells treated with increasing concentrations of the dynein inhibitor Ciliobrevin D for the indicated times prior to infection with C5/D64-HVR1. In A, the image was obtained at a resolution of 50 μm, and grayscale intensity correlates with eGFP expression. In B, data are normalized to infection of untreated control cells. Data are the means ± SD of at least 3 independent replicates. (C) WT and CAR KO A549 cells were infected with either the C5/D64-HVR1 chimera virus or EV-A71 control virus with or without 5 µM HD5. Where indicated, cells were pretreated with 67 μM Ciliobrevin D for 4 h prior to infection, and the inhibitor remained present for an additional 2 h p.i. Each point is an independent replicate (n > 3), and bars are the means ± SD of the percent infectivity compared to untreated control cells. Data were analyzed by ordinary two-way ANOVA with Šídák’s multiple comparisons test with a single pooled variance comparing infection with and without Ciliobrevin D for each virus. ****, *P* < 0.0001; ns = *P* > 0.05. Representative wells of a 96-well plate are shown below, where grayscale intensity correlates with eGFP expression (C5/D64-HVR1) or immunofluorescence (EV-A71), indicative of successful infection.

### HAdV traffics to the MTOC en route to the nucleus upon α-defensin-mediated infection

Following microtubule-mediated transport, HAdV reaches the MTOC. In canonical HAdV entry, the perinuclear MTOC serves as a transition point for disengagement from microtubules, enabling HAdV to bind to the NPC [37,38]. To determine whether the α-defensin-mediated entry route also converges on the MTOC prior to nuclear entry, we used Leptomycin B (LMB). Although the exact mechanism is unknown, LMB prevents HAdV disengagement from the MTOC and blocks infection by limiting the availability of one or more cellular factors that are in turn dependent on nuclear export mediated by chromosomal maintenance 1 (CRM1) [6,39]. We first treated cells with varying concentrations of LMB for 30 min and assessed C5/D64-HVR1 infection by the classical pathway, the HD5-mediated pathway, or when both pathways were available. As a control, we also included MA104 cells infected with rhesus rotavirus (RRV), which is an RNA virus that replicates in the cytoplasm and does not require MTOC-directed transport for genome delivery [40]. We found that C5/D64-HVR1 infection in all groups was almost completely inhibited by 20 nM LMB (Fig 4A-B). RRV infection remained unaffected, confirming the specificity of MTOC disruption. To broaden these findings, we also tested HNP1, a myeloid α-defensin that we previously showed could also rescue C5/D64-HVR1 infection of CAR KO A549 cells [23], and observed similar results (Fig 4C-D).

**Fig 4.**
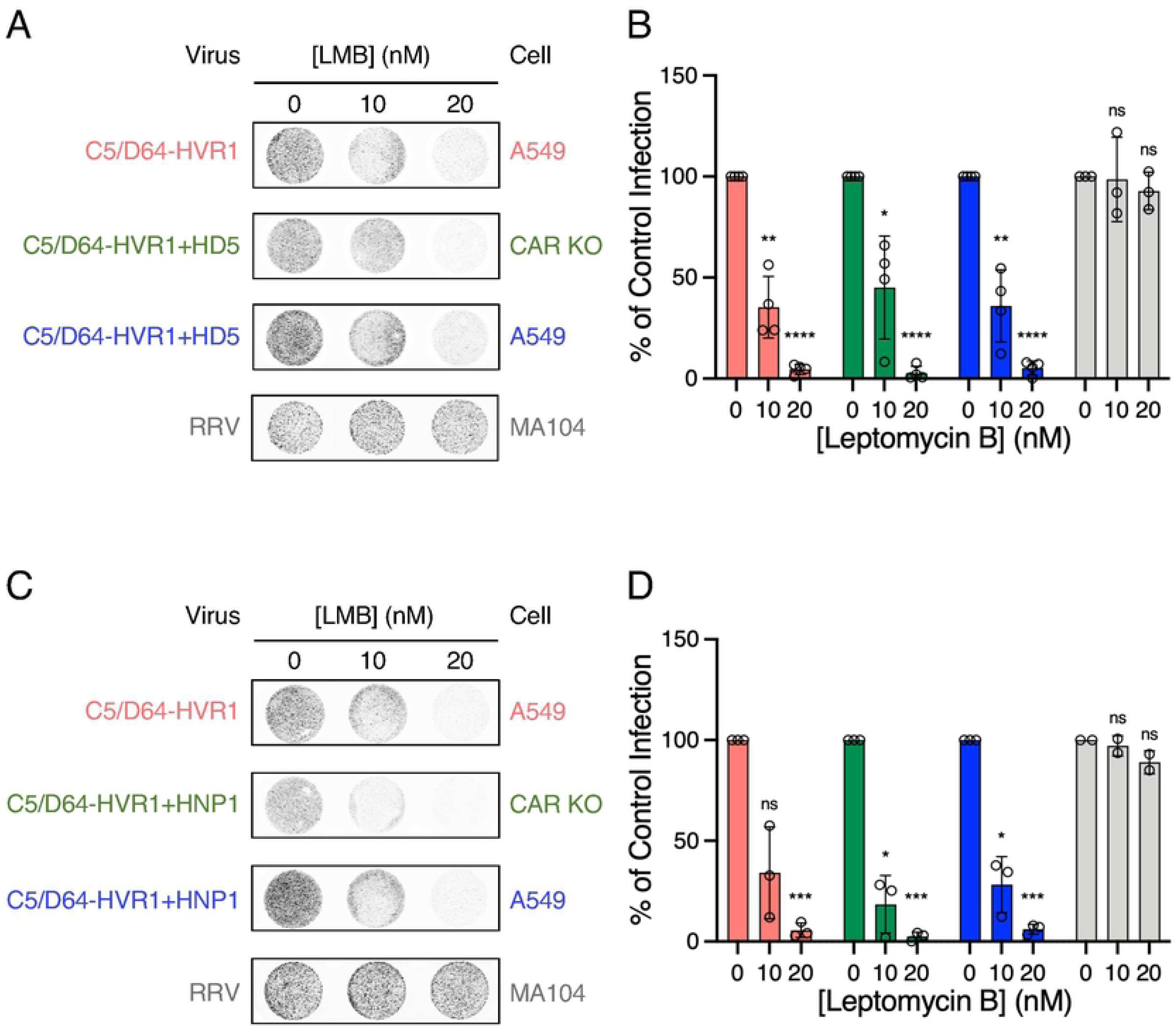
α-Defensin-mediated HAdV entry is CRM1-dependent. (A, C) Representative images and (B, D) quantification of infection of WT A549, CAR KO A549, or MA104 cells infected with either the C5/D64-HVR1 chimera virus or RRV control virus with or without (A, B) 5 µM HD5 or (C, D) 5 µM HNP1. Cells were pretreated or not with the indicated concentrations of the CRM1 inhibitor leptomycin B (LMB) for 30 min prior to infection. Images in A and C were obtained at a resolution of 50 μm, and grayscale intensity correlates with eGFP expression (C5/D64-HVR1) or immunofluorescence (RRV). In B and D, colors correspond to the conditions in panels A and C, each point is an independent replicate (n > 3), and bars are the means ± SD of the percent infectivity compared to untreated control cells. Data were analyzed by repeated measures two-way ANOVA with the Geisser-Greenhouse correction comparing LMB-treated samples with control, untreated samples for each virus using Dunnett’s multiple comparisons test with individual variances computed for each comparison. *, *P* = 0.01 to 0.05; **, *P* = 0.001 to 0.01; ***, P = 0.0001 to 0.001; ****, *P* < 0.0001.; ns = *P* > 0.05.

To directly visualize whether MTOC disruption affects viral nuclear entry, we performed immunofluorescence staining. We infected cells with AF488-labeled C5/D64-HVR1 in the presence or absence of 20 nM LMB and assessed nuclear localization. By 60 min p.i., approximately 80% of C5/D64-HVR1 was in the nucleus via both pathways (Fig 5A and C). Consistent with its effect on infection, LMB treatment significantly reduced this proportion to approximately 40% for both pathways. Similar results were obtained with HNP1 (Fig 5B and D). Collectively, these results demonstrate that HAdV traffics through the MTOC upon α-defensin-mediated entry. Thus, despite binding to cells independently of canonical primary receptors, HAdV that enters cells via α-defensin binding follows a path like that of conventional infection.

**Fig 5.**
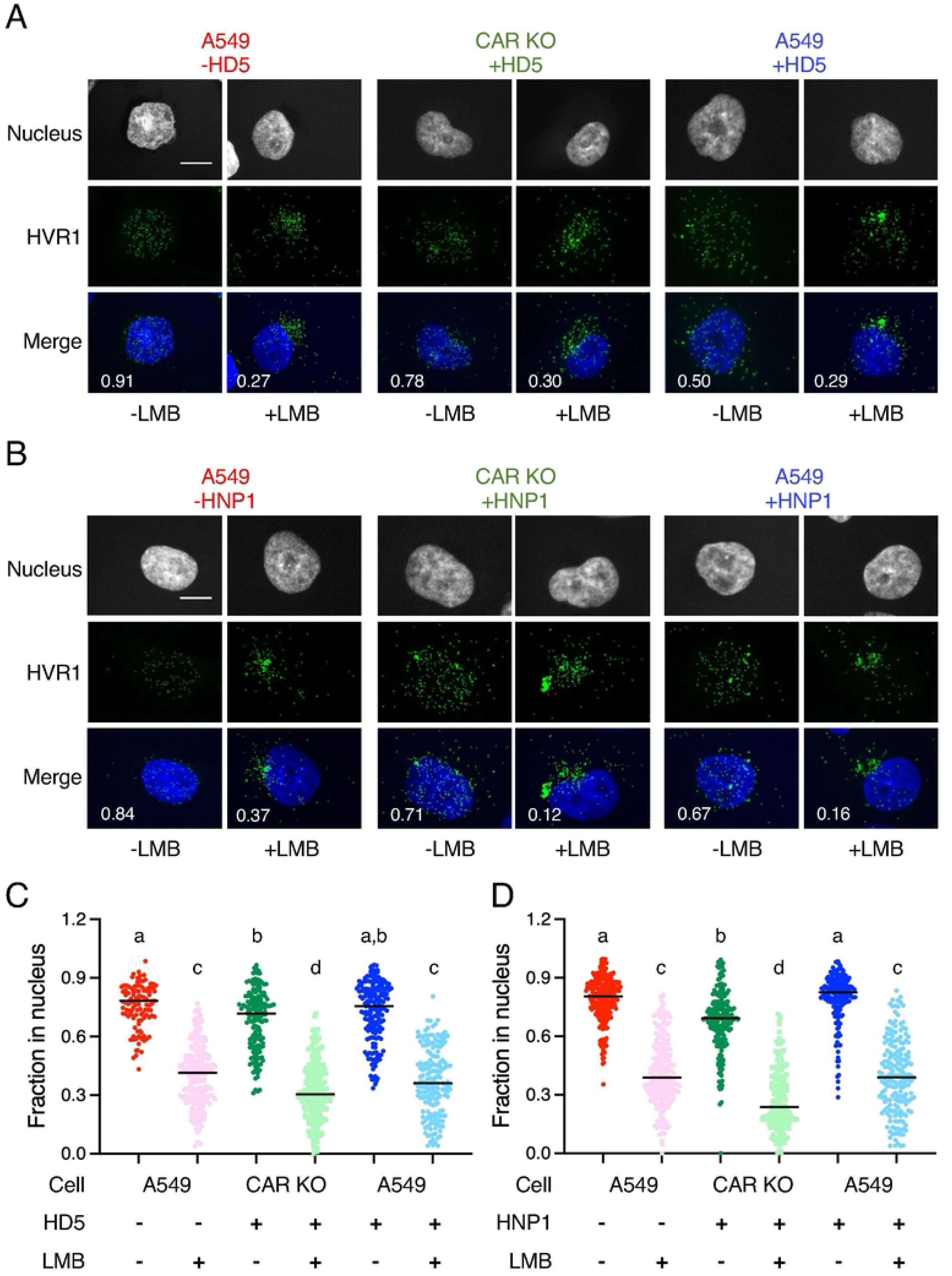
CRM1 inhibition traps HAdV at the microtubule organizing center upon canonical and α-defensin-mediated entry. Representative images of WT or CAR KO A549 cells infected with Alex Fluor 488 (AF488)-labeled C5/D64-HVR1 in the presence or absence of (A) 5 µM HD5 or (B) 5 µM HNP1. Cells were pretreated or not with 20 nM CRM1 inhibitor, leptomycin B (LMB), for 30 min prior to infection. Images are maximum intensity Z-projections from 60 min or 90 min p.i. Note that LMB was removed from cells during the infection period. Merged images include C5/D64-HVR1 (AF488, green) and the nucleus (DAPI, blue). Scale bars indicate 10 μm. The fraction of AF488 signal in the nucleus is indicated for each merged image. (C, D) The fraction of AF488-labelled C5/D64-HVR1 in the nucleus was assessed for the indicated combinations of cells, defensins, and LMB by confocal microscopy and automated image analysis. Each point is an individual cell (n>70) from one of three biological replicates that were pooled and analyzed in aggregate. Horizontal bars indicate means. Data were analyzed by ordinary one-way ANOVA with Tukey’s multiple comparisons test with a single pooled variance comparing the mean of each column with every other column. Different letters indicate significant differences between groups (*P* < 0.05). Groups that share a letter are not statistically significantly different from one another.

### HAdVs retain their intrinsic defensin sensitivity upon co-infection

The equivalence of intracellular trafficking via canonical and defensin-mediated routes suggests that altered intracellular trafficking does not underlie the defensin-resistant phenotypes of engineered (e.g., C5/D64-HVR1) or naturally occurring (e.g., HAdV-D64) serotypes, which instead reflects capsid-intrinsic properties. This model predicts that viruses will maintain their phenotypes even during co-infection. To test this, WT A549 cells were infected with a mixture of HAdV-C5 (mCherry) and HAdV-D64 (eGFP) in the presence of increasing HD5 concentrations. Consistent with our hypothesis, HAdV-D64 remained resistant while HAdV-C5 was neutralized (Fig 6A). Control infections with two HAdV-C5 strains expressing different fluorophores confirmed that both were equivalently neutralized, ruling out effects of reporter or mixed infection (Fig 6B). To directly assess infection at the single-cell level, cells were co-infected with Cy3-labeled HAdV-C5 and AF488-labeled C5/D64-HVR1, and nuclear colocalization was quantified at 90 min p.i. In the absence of HD5, 75% of HAdV-C5 and 85% of C5/D64-HVR1 capsids reached the nucleus (Fig 6C and 6D). This decreased to 34% and 45%, respectively, in the presence of 10 µM HD5. Similar values were observed in single infections (31% vs. 51%), indicating that each virus retains its intrinsic phenotype within the same cell and that defensin sensitivity is not altered by co-infection.

**Fig 6.**
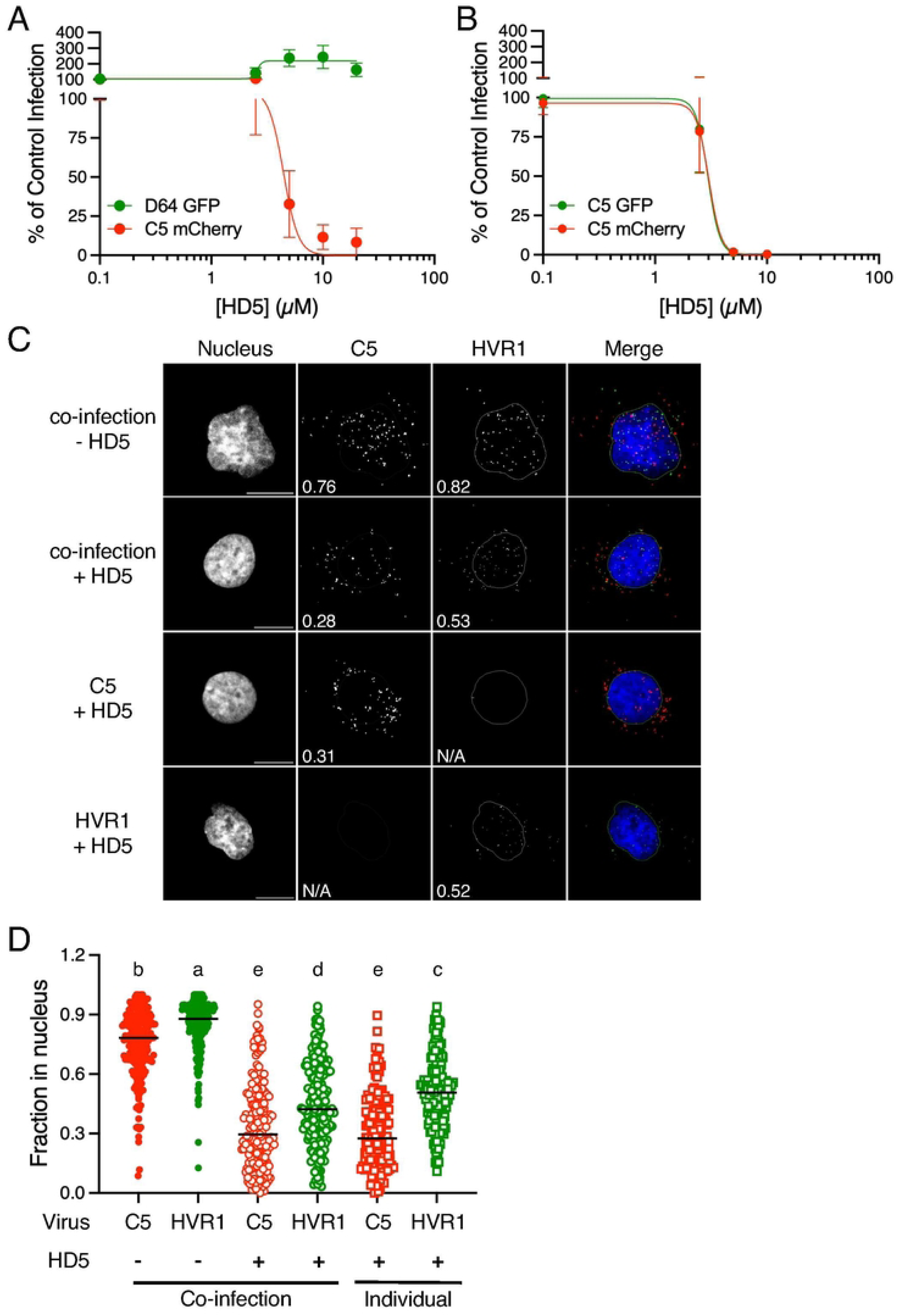
HAdVs retain their intrinsic defensin sensitivity upon co-infection. (A, B) Quantification of infection of WT A549 cells co-infected with the indicated viruses in the presence of increasing concentrations of HD5. Data are normalized to infection in the absence of HD5 for each virus and are the means ± SD of 3 independent replicates. C) Representative images of WT A549 cells co-infected or singly infected with Cy3-labeled HAdV-C5 or Alex Fluor 488 (AF488)-labeled C5/D64-HVR1 in the presence or absence of 10 µM HD5. Images are maximum intensity Z-projections from 90 min p.i. Merged images include C5/D64-HVR1 (AF488, green), HAdV-C5 (Cy3, red), and the nucleus (DAPI, blue). Lines depict nuclear boundaries used for quantitation. Scale bars indicate 10 μm. The fraction of virus colocalized with the nucleus is indicated for each image. (D) The fraction of each virus in the nucleus was assessed by confocal microscopy and automated image analysis. Each point is an individual cell (n>166) from one of three biological replicates that were pooled and analyzed in aggregate. Horizontal bars indicate means. Data were analyzed by ordinary one-way ANOVA with Tukey’s multiple comparisons test with a single pooled variance comparing the mean of each column with every other column. Different letters indicate significant differences between groups (*P* < 0.05). Groups that share a letter are not statistically significantly different from one another.

## Discussion

This study resolves a central paradox in adenovirus/defensin biology: α-defensins act as both broad antiviral restriction factors and, for resistant capsids, proviral attachment factors. This duality raises a basic question—does enhanced infection reflect access to a distinct, defensin-permissive trafficking pathway, or does it operate within the same conserved intracellular route used by canonical entry? Our data support the latter and indicate that for α-defensin-resistant HAdVs, α-defensins facilitate infection not by influencing an intracellular step but by acting extracellularly as molecular bridges that mediate initial attachment. Once internalized following attachment via canonical viral receptors or α-defensin-mediated binding, virions follow the same pathway: endocytic uptake, protein VI-dependent endosomal escape, dynein-driven microtubule transport, perinuclear accumulation at the MTOC, and NPC engagement (Fig 7). Accordingly, as we previously speculated [23], susceptibility to α-defensin-mediated neutralization is determined by intrinsic capsid features within this common entry pathway rather than by diversion of sensitive and resistant virions into distinct trafficking routes. These findings highlight the essentiality of the HAdV intracellular route: receptor usage is comparatively flexible, whereas post-entry pathways are constrained.

**Fig 7.**
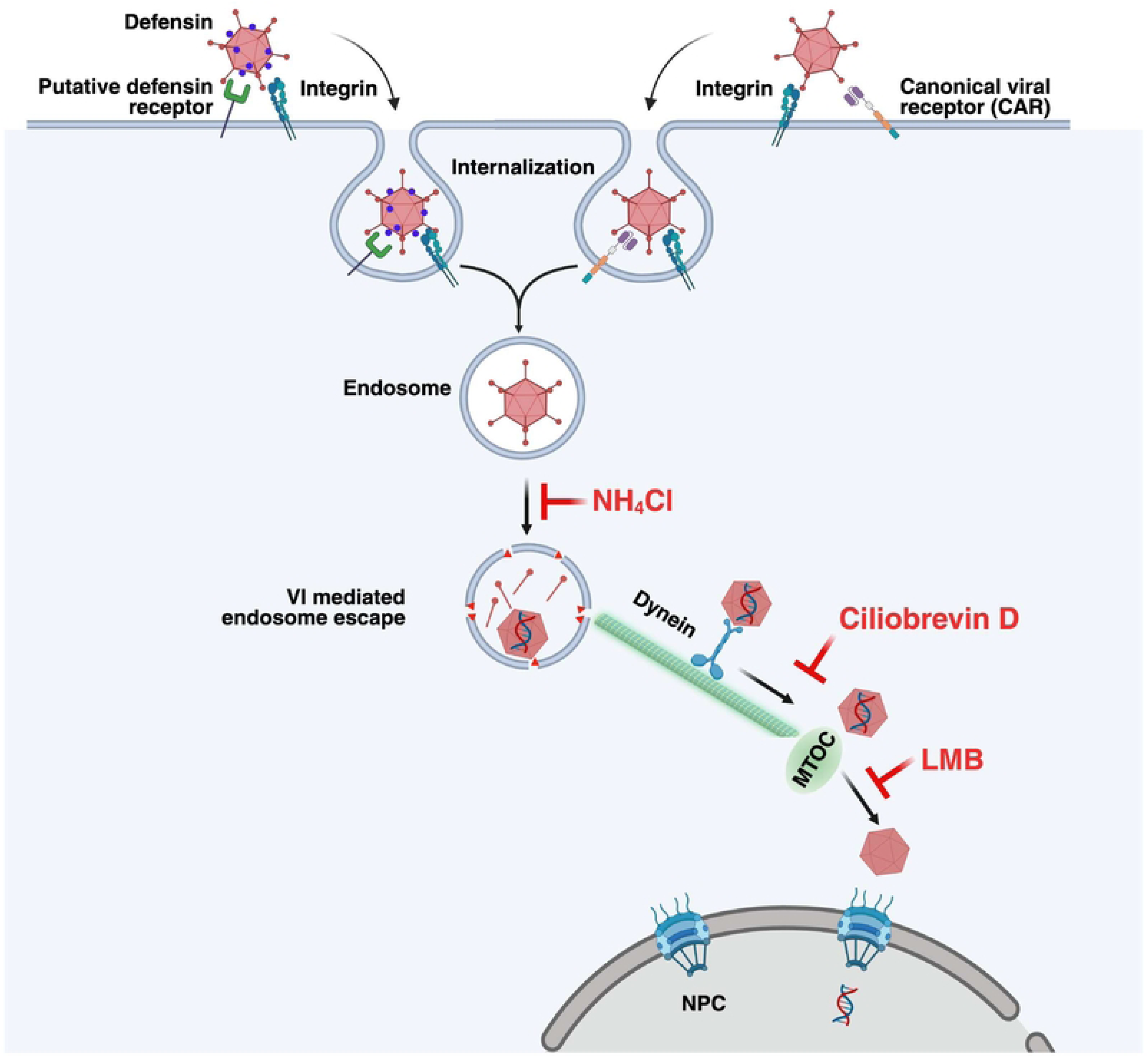
Model of HAdV intracellular trafficking. Schematic representation of HAdV infection upon attachment to cells via α-defensin-mediated binding or canonical viral receptors (e.g., CAR). Attachment via either mechanism converges on a common trafficking route whereby the virus is internalized, escapes from endosomes, and co-opts host cytoplasmic machinery for directed transport toward the nucleus. Key host components include a putative α-defensin receptor at the cell surface, dynein motor proteins for microtubule-based transport, the microtubule-organizing center (MTOC), and the nuclear pore complex (NPC). Blue dots represent α-defensins bound to the viral capsid. Black arrows indicate the direction of viral movement. Inhibitors used in this study are indicated in red. Figure created with BioRender.com.

A key implication of our study is that α-defensins internalized with HAdV do not provide membrane lytic activity in endosomes. Even though HD5 has membrane lytic activity [41–43], in the context of HAdV entry it fails to substitute for protein VI. One possible explanation is that HD5 remains bound to the viral capsid during endosomal transit, limiting the pool of free peptide available for membrane interaction. Alternatively, the endosomal environment or lipid composition of the endosomal membrane may not favor HD5-mediated membrane permeabilization. Biophysical studies have shown that α-defensin membrane activity depends critically on lipid composition [44–46]. A third explanation is that membrane perturbations mediated by HD5 are insufficient for adenovirus endosome escape. Unlike α-defensins that form small pores [47,48], protein VI induces catastrophic membrane fragmentation, reducing liposomes to small vesicles and enabling translocation of large molecules [33,49]. These mechanisms are not mutually exclusive but collectively indicate that HD5 cannot replace the membrane-disrupting activity of protein VI. Hence, productive endosomal escape still requires the precise temporal and spatial exposure of protein VI during regulated viral uncoating.

Since our results suggest that α-defensins act solely by mediating attachment, a key question is how the α-defensin engagement with its putative receptor can functionally substitute for fiber binding to a canonical receptor such as CAR. One biophysical model for the early events in HAdV entry from Greber and colleagues proposes a CAR/integrin ‘tug-of-war’ at the plasma membrane that generates mechanical force at the plasma membrane—CAR/fiber interactions permit lateral drift while integrin/PB binding spatially confines the virus, together driving fiber shedding and activation of protein VI [50]. However, productive infection still occurs upon cell attachment through bridging molecules like α-defensins and blood factors that bind to sites on the capsid other than the fiber knob. This suggests either that these interactions mediate sufficient drifting motions to allow for the integrin-dependent “tug of war” or that mechanical uncoating at the plasma membrane can be bypassed in favor of uncoating in endosomes in response to local cues. Thus, a requirement for integrin binding by PB for productive infection may primarily reflect a need for integrin signaling and correct endosomal routing rather than integrin anchoring, underscoring that α-defensin-mediated attachment feeds into a conserved, integrin-dependent entry program rather than creating a fundamentally distinct pathway.

These mechanistic insights have biological and translational implications. In defensin-rich milieus, such as neutrophil-infiltrated airways during acute respiratory infections or the intestinal lumen into which Paneth cells secrete HD5, defensin-resistant HAdVs may exhibit expanded tropism by attaching to cells that lack canonical primary receptors. For vector development, this has immediate practical implications. Defensin resistant, HAdV-C5-based oncolytic vectors could show enhanced tumor penetration in inflammatory microenvironments where defensins are elevated. Similarly, gene therapy vectors derived from α-defensin-resistant serotypes might exhibit off-target transduction in defensin-rich tissues such as inflamed joints in arthritis patients or inflammatory bowel disease lesions where both HD5 and HNPs are abundant. Conversely, intentionally engineering defensin binding capacity while maintaining resistance to the uncoating block could enhance vector delivery to specific tissues. Nevertheless, like HAdVs that enter via canonical receptors, tropism may be limited at downstream trafficking and replication steps.

We acknowledge the limitations of our interpretations. The earliest capsid-uncoating events during defensin-mediated entry are inferred rather than directly visualized, leaving the precise molecular trigger for endosomal disassembly unresolved. In addition, these experiments were performed in tumor cell lines, which may not fully reflect receptor expression patterns or α-defensin exposure in tissues. And the work focuses exclusively on a CAR-utilizing adenovirus; however, our prior work also included studies of a HAdV that uses CD46 as a primary receptor [23]. Although pharmacological perturbations, genetic mutants, and imaging collectively support the conclusion that both entry routes converge on conserved intracellular checkpoints, these approaches cannot fully exclude off-target effects or subtle mechanistic differences in integrin signaling or trafficking adaptors. Extending the analysis to other HAdV species and serotypes, to untransformed human cells, and *in vivo* will be essential to defining the generality and physiological relevance of the mechanisms proposed here.

In summary, α-defensins enhance infection of defensin-resistant HAdVs mainly by increasing attachment, while intracellular trafficking proceeds along a route common to receptor-mediated entry. Defensin resistance is therefore determined by capsid features that dictate how α-defensins alter uncoating rather than by rerouting resistant viruses into a distinct, defensin-insensitive endocytic pathway or by having α-defensins substitute for the membrane lytic activity of protein VI. This framework clarifies the dual, serotype-dependent antiviral and proviral nature of α-defensins within the constraints of a conserved entry pathway inside the cell.

## Materials and Methods

### Cells

A549 (ATCC), 293β5 [13], RD (a kind gift from Michael Gale, Jr. of the University of Minnesota, Minneapolis, MN), and MA104 (a kind gift from Monica McNeal of Cincinnati Children’s Hospital Medical Center, Cincinnati, OH) cells were maintained in Dulbecco’s modified Eagle’s medium (DMEM) with 10% fetal bovine serum (FBS), penicillin, streptomycin, L-glutamine, and non-essential amino acids (complete DMEM). Clonal CAR KO A549 cells were generated as previously described and cultured in puromycin (1.5 µg/mL) containing medium [23]. All cells were tested at least quarterly for mycoplasma contamination.

### Viruses

The HAdV-C5-base vectors used in these studies are replication-defective, E1/E3-deleted, and contain a CMV promoter-driven enhanced green fluorescent protein (CMV-eGFP) reporter gene cassette in place of E1. Construction of C5/D64-HVR1 was previously described [11,23]. The protease P137L mutant was generated by recombineering the BACmid containing the C5/D64-HVR1 genome to mutate the codon for proline 137 in the *L3/23K* gene (CCC) to leucine (CTC). The C5/D64-HVR1-VI L40Q mutant was also generated by recombineering the BACmid containing the C5/D64-HVR1 genome to mutate the codon for leucine 40 in the *protein VI* gene (CTG) to glutamine (CAA). The fidelity of all BACmid constructs was verified by Sanger sequencing of the recombineered region and by restriction digest of the entire BACmid. HAdV-D64 expressing eGFP was previously described [51]. The HAdV-C5 vector expressing mCherry was obtained from Vector Biolabs (Cat: 1796). All AdV vectors were propagated in 293β5 cells at 37°C, except the protease P137L mutant, which was propagated at the non-permissive temperature (39.5°C), as described previously [33]. Virions were purified by CsCl-gradient centrifugation; dialyzed against 150 mM NaCl, 40 mM Tris, 10% glycerol, 2 mM MgCl_2_, pH 8.1; flash frozen in liquid nitrogen; stored at −80°C; and quantified as previously described [11,14]. For microscopy, viruses were labeled with Alexa Fluor 488 (AF488) carboxylic acid, tetrafluorophenyl ester (Thermo Fisher Scientific) or Cy3 monoreactive dye (GE Healthcare) as previously described [15,52].

Enterovirus A71 (EV-A71) strain USA/2018-20932 was obtained from BEI Resources (Cat: NR-52000; Lot: 70032015) and propagated on RD cells. The original stock was diluted to 1.3 x 10^6^ PFU/mL in serum-free media (SFM) and used to infect a T-25 flask of RD cells at 80-90% confluency. After 1 h, the inoculum was removed and replaced with DMEM supplemented with 2.5% FBS. The culture was harvested 4 d post-infection (p.i.), upon the appearance of complete cytopathic effect. Cells and supernatant were separated. Cells were lysed by three freeze/thaw cycles and centrifuged to remove debris. Clarified lysate and supernatant were then combined, flash frozen in liquid nitrogen, and stored at −80°C. This P1 stock was amplified by two additional passages on RD cells prior to use in experiments.

Rhesus rotavirus (RRV) was a kind gift from Monica McNeal of Cincinnati Children’s Hospital Medical Center (Cincinnati, OH, USA). RRV was amplified on MA104 cells, and viral lysate was prepared as previously described [17].

### α-Defensin peptides

HNP1 and HD5 were created using peptides synthesized by CPC Scientific (Sunnyvale, CA) or LifeTein (Somerset, NJ), which were oxidatively re-folded, purified by reverse-phase high-pressure liquid chromatography (RP-HPLC) to homogeneity, lyophilized, resuspended in deionized water, and quantified by absorbance at 280 nm as described [11,17]. Purity (>99%) and mass were verified by analytical RP-HPLC and mass spectrometry.

### Quantification of viral infection

To assess the effects of ammonium chloride (NH₄Cl) on viral infectivity, confluent monolayers of WT or CAR KO A549 cells in black wall, clear bottom 96-well plates (PerkinElmer, San Jose, CA) were washed once with RT SFM, and 50 µL of either 50 mM NH₄Cl or SFM was added. Plates were returned to the incubator for 1 h at 37°C. Purified C5/D64-HVR1 was incubated with or without 5 µM HD5 for 45 min on ice in SFM, after which 50 µL of the C5/D64-HVR1/HD5 mixture, C5/D64-HVR1 alone, or EV-A71 P3 alone was added to the cells, yielding a final volume of 100 µL/well and a concentration of each virus that was predetermined to yield 50-70% of maximal signal in the absence of inhibitors. Following a 2 h incubation at 37°C, cells were washed with complete DMEM and cultured with 100 µL of complete DMEM (with or without 50 mM NH₄Cl) at 37°C for 8 h. EV-A71 samples were fixed and stained with a pan-enterovirus monoclonal antibody (clone L66J; Invitrogen, PIMA518206) at a 1:400 dilution for 1 h at RT, followed by Alexa Fluor 488-conjugated goat anti-mouse IgG (Invitrogen) at a 1:1000 dilution for 45 min at RT. HAdV samples were washed with phosphate-buffered saline (PBS). Plates were scanned for AF488/eGFP signal using a Sapphire (Azure) imager. For all samples, Fiji (version 2.1.0/1.53c) was used to quantify background-subtracted total monolayer fluorescence, and data are shown as a percent of control infection in the absence of NH₄Cl.

To investigate a possible role for HD5 in endosome escape, C5/D64-HVR1 and pVI L40Q mutant virus stocks were normalized by particle number based on a Bio-Rad Protein Assay, and an equivalent serial dilution of each virus was used to infect A549 cells. To assess the effects of HD5, CAR KO A549 cells were infected with the same particle-based multiplicity of infection (MOI) (∼4200 particles/cell) of C5/D64-HVR1 and pVI L40Q mutant viruses that had been incubated with or without HD5 for 45 min on ice. For all experiments, infected cells were incubated for 20-24 h, washed with PBS, scanned, and quantified as above.

To analyze intracellular virus trafficking, the cytoplasmic dynein inhibitor Ciliobrevin D was used to block microtubule-dependent transport. To determine the optimal Ciliobrevin D concentration, A549 cells were treated with 50 µL of SFM with or without increasing concentrations of Ciliobrevin D and incubated for 1, 2, 3, or 4 h at 37°C. An equivalent volume of C5/D64-HVR1 virus was then added, and samples were incubated for an additional 1.5 h at 37°C. Virus and inhibitor were removed, and cells were incubated for 20-24 h, washed with PBS, and scanned for eGFP signal as above. Based on these results, treatment with 67 µM Ciliobrevin D for 4 h was used for subsequent experiments. Purified C5/D64-HVR1 virus was incubated with or without 5 µM HD5 for 45 min on ice in SFM. Either 50 µL of the C5/D64-HVR1/HD5 mixture, C5/D64-HVR1 virus alone, or EV-A71 P3 alone was added to inhibitor-pretreated WT or CAR KO A549 cells. Cells were incubated at 37°C for 2 h to permit viral entry, after which the inoculum was removed and replaced with complete DMEM. Cells were cultured for an additional 20-24 h, washed with PBS (HAdV) or fixed and stained for immunofluorescence (EV-A71), scanned, and quantified as above.

To assess the accumulation of virus at the microtubule-organizing center (MTOC), leptomycin B (LMB) was used. A549, CAR KO A549, or MA104 cells were treated with 50 µL SFM with or without 10 or 20 nM LMB for 30 min on ice. Purified C5/D64-HVR1 was incubated with or without 5 µM HD5 or HNP1 for 45 min. RRV was activated with 10 μg/mL type IX-S EDTA-free porcine trypsin for 1 h at 37°C. Following removal of LMB by washing, 50 μL C5/D64-HVR1/defensin mixture or C5/D64-HVR1 alone was added to A549 or CAR KO A549 cells and activated RRV alone was added to MA104 cells. Following a 2 h incubation at 37°C, the inoculum was removed and cells were cultured with 100 µL of complete DMEM at 37°C for 20-24 h. RRV samples were fixed and immunostained with a rabbit anti-rotavirus primary antibody followed by Alexa Fluor 488-conjugated goat anti-rabbit secondary antibody (Invitrogen) as previously described [17]. HAdV samples were washed with PBS. Plates were scanned for AF488/eGFP signal using a Sapphire imager and quantified as above.

To assess the effects of HD5 on co-infection, purified HAdV-C5 expressing eGFP, HAdV-C5 expressing mCherry, or HAdV-D64 expressing eGFP was incubated with or without increasing concentrations of HD5 for 45 min on ice in SFM. Samples (50 µL/well) were added to WT A549 cells that had been washed twice with SFM and incubated at 37°C for 2 h. The inoculum was removed, cells were washed once with SFM, and cells were cultured at 37°C for 20-24 h in complete, phenol red-free DMEM before being scanned on a Typhoon 9400 (GE Healthcare) and quantified as above.

### SDS-PAGE analysis of the C5/D64-HVR1 protease P137L mutant

Equal amounts (250 ng) of HAdV-C5 and the C5/D64-HVR1 protease P137L mutant were diluted in 5x SDS loading buffer [3.2% SDS, 100 mM Tris, 0.04% bromophenol blue, 16% glycerol (w/v), 200 mM mercaptoethanol, pH 6.8]. Samples were heated at 100°C for 10 min, cooled to RT, briefly centrifuged, and loaded (30 µl) onto a 12% polyacrylamide gel. Electrophoresis was performed in Tris-glycine-SDS running buffer (25 mM Tris, 192 mM glycine, 0.1% SDS) at 30 mA for 45 min or until the dye front approached the gel bottom. Gels were stained with AzureRed Fluorescent Total Protein Stain (Azure Biosystems) according to the manufacturer’s instructions, rinsed in deionized water, and imaged on a Sapphire imager.

### Endosomal colocalization assay

WT or CAR KO A549 cells were sparsely seeded on glass coverslips and cultured overnight. Alexa Fluor 488-labeled C5/D64-HVR1 or protease P137L mutant viruses were incubated with or without 5 µM HD5 for 45 min on ice in SFM. Cells were washed twice with cold SFM, and 100 µL of the virus/defensin mixture or virus alone was added per coverslip. After incubation for 45 min on ice, cells were washed twice with RT SFM and incubated with 100 µL of RT SFM at 37°C for 30 min to allow viral entry. Coverslips were then washed with PBS and fixed in 2% paraformaldehyde (PFA) in PBS for 15 min at RT. Fixed cells were washed once with PBS and permeabilized for 20 min at RT using permeabilization buffer (20 mM glycine, 0.5% Triton X-100 in PBS). Cells were sequentially stained with mouse monoclonal anti-EEA1 antibody (clone 14, BD Biosciences: AB_397830, 1:200) diluted in Pierce Immunostain Enhancer for 1 h at RT, followed by Alexa Fluor 555-conjugated goat anti-mouse IgG secondary antibody (Invitrogen; 1:500) diluted in Immunostain Enhancer for 45 min at RT. Nuclei were counterstained with 500 ng/mL DAPI for 5 min. Coverslips were mounted using ProLong Gold Antifade Mountant (Life Technologies: P36930).

### Nuclear entry assay

WT or CAR KO A549 cells were pretreated with or without 20 nM LMB for 30 min on ice before infection. Alexa Fluor 488-labeled C5/D64-HVR1 was incubated with or without 5 µM HD5 or HNP1 for 75 min on ice in SFM. Media was removed and 100 µL of the virus/defensin mixture or virus alone was added per coverslip. After incubation for 45 min on ice, coverslips were washed with SFM and incubated at 37°C for 60-90 min to allow viral entry. Coverslips were then fixed, permeabilized, stained with DAPI, and mounted as above.

For co-infection assay and controls, Alexa Fluor 488-labeled C5/D64-HVR1 and/or Cy3-labeled HAdV-D64 was incubated with WT A549 cells in SFM for 45 min at 4°C. Unbound virus was removed by washing with ice cold SFM. Samples were incubated with or without 10 µM HD5 in SFM for 45 min at 4°C then shifted to 37°C for 90 min to allow viral entry. Coverslips were washed, fixed, permeabilized, stained with DAPI, and mounted as above.

### Image acquisition and analysis

Z-stack images covering the entire cell volume were acquired using a Zeiss LSM 800 confocal laser-scanning microscope equipped with a 100× objective. Maximum intensity z-projections of each color channel for each field of view were generated using Fiji (v2.14.0/1.54h). Background thresholds for Alexa Fluor 488-labeled virus signal and EEA1 were defined manually for each replicate in Fiji using uninfected negative controls. A human-in-the-loop model was trained in CellPose 2.0 to identify cell borders and generate cell masks [53]. CellPose masks and z-projection images were imported into CellProfiler (v4.2.5) [54], and background-subtracted single-cell images were identified and exported. A second CellProfiler pipeline was then used to analyze the single-cell images. Nuclei were identified using DAPI by thresholding using the Otsu method in CellProfiler or via CellPose 2.0. For endosomal colocalization, Manders’ coefficient was used to quantify the integrated Alexa Fluor 488 intensity within EEA1-positive regions per cell. For nuclear entry analysis, Alexa Fluor 488 intensity within the whole cell and the nuclear region were measured on a per-cell basis.

### Statistical analysis

Statistical tests were performed using Prism 11.0.2 as indicated in the figure legends.

## Acknowledgements

The authors thank Kaitlin Hulce for her assistance in propagating EV-A71. This work was supported by R01 AI104920, from the National Institute of Allergy and Infectious Diseases (www.niaid.nih.gov) and by the Office of the Director, National Institutes of Health (www.nih.gov/institutes-nih/nih-office-director) under Award Number S10 OD026741. Additional support in the form of subsidized core services was provided by NIH grants UL1 TR000423 from the National Center for Advancing Translational Sciences (ncats.nih.gov), and P30 CA015704 from the National Cancer Institute (www.cancer.gov). The funders had no role in study design, data collection and analysis, decision to publish, or preparation of the manuscript.

## Notes

### Competing Interest Statement

The authors have declared no competing interest.

## References

1. Lion T. Adenovirus persistence, reactivation, and clinical management. FEBS Lett. 2019;593: 3571–3582. doi:10.1002/1873-3468.13576

2. MacNeil KM, Dodge MJ, Evans AM, Tessier TM, Weinberg JB, Mymryk JS. Adenoviruses in medicine: innocuous pathogen, predator, or partner. Trends Mol Med. 2023;29: 4–19. doi:10.1016/j.molmed.2022.10.001

3. Kajon AE. Adenovirus infections: new insights for the clinical laboratory. J Clin Microbiol. 2024;62: e0083622. doi:10.1128/jcm.00836-22

4. Stasiak AC, Stehle T. Human adenovirus binding to host cell receptors: a structural view. Med Microbiol Immunol. 2019. doi:10.1007/s00430-019-00645-2

5. Greber UF, Flatt JW. Adenovirus Entry: From Infection to Immunity. Annu Rev Virol. 2019. doi:10.1146/annurev-virology-092818-015550

6. Greber UF, Suomalainen M. Adenovirus Entry - Stability, Uncoating and Nuclear Import. Mol Microbiol. 2022. doi:10.1111/mmi.14909

7. Moyer CL, Wiethoff CM, Maier O, Smith JG, Nemerow GR. Functional genetic and biophysical analyses of membrane disruption by human adenovirus. J Virol. 2011;85: 2631–41. doi:10.1128/JVI.02321-10

8. Charman M, Herrmann C, Weitzman MD. Viral and cellular interactions during adenovirus DNA replication. FEBS Lett. 2019;593: 3531–3550. doi:10.1002/1873-3468.13695

9. Tartaglia LJ, Badamchi-Zadeh A, Abbink P, Blass E, Aid M, Gebre MS, et al. Alpha-defensin 5 differentially modulates adenovirus vaccine vectors from different serotypes in vivo. PLoS Pathog. 2019;15: e1008180. doi:10.1371/journal.ppat.1008180

10. Zhao C, Porter JM, Burke PC, Arnberg N, Smith JG. Alpha-defensin binding expands human adenovirus tropism. Dryad Digit Repos. 2024. doi:10.5061/dryad.d2547d89h

11. Diaz K, Hu CT, Sul Y, Bromme BA, Myers ND, Skorohodova KV, et al. Defensin-driven viral evolution. PLoS Pathog. 2020;16: e1009018. doi:10.1371/journal.ppat.1009018

12. Wilson SS, Bromme BA, Holly MK, Wiens ME, Gounder AP, Sul Y, et al. Alpha-defensin-dependent enhancement of enteric viral infection. PLoS Pathog. 2017;13: e1006446. doi:10.1371/journal.ppat.1006446

13. Smith JG, Silvestry M, Lindert S, Lu W, Nemerow GR, Stewart PL. Insight into the mechanisms of adenovirus capsid disassembly from studies of defensin neutralization. PLoS Pathog. 2010;6: e1000959. doi:10.1371/journal.ppat.1000959

14. Nguyen EK, Nemerow GR, Smith JG. Direct evidence from single-cell analysis that human alpha-defensins block adenovirus uncoating to neutralize infection. J Virol. 2010;84: 4041–9. doi:10.1128/JVI.02471-09

15. Smith JG, Nemerow GR. Mechanism of adenovirus neutralization by Human alpha-defensins. Cell Host Microbe. 2008;3: 11–9. doi:10.1016/j.chom.2007.12.001

16. Porter JM, Hulce KR, Oswald MS, Busuttil K, Emmanuel SN, Bennett A, et al. Mechanisms of AAV neutralization by human alpha-defensins. PLOS Pathog. 2025;21: e1013283. doi:10.1371/journal.ppat.1013283

17. Hu CT, Diaz K, Yang LC, Sharma A, Greenberg HB, Smith JG. Corrected and republished from: “VP4 Is a Determinant of Alpha-Defensin Modulation of Rotaviral Infection.” J Virol. 2023;97: e0096223. doi:10.1128/jvi.00962-23

18. Porter JM, Oswald MS, Sharma A, Emmanuel S, Kansol A, Bennett A, et al. A Single Surface-Exposed Amino Acid Determines Differential Neutralization of AAV1 and AAV6 by Human Alpha-Defensins. J Virol. 2023;97: e0006023. doi:10.1128/jvi.00060-23

19. Holly MK, Diaz K, Smith JG. Defensins in Viral Infection and Pathogenesis. Annu Rev Virol. 2017;4: 369–391. doi:10.1146/annurev-virology-101416-041734

20. Wilson SS, Wiens ME, Smith JG. Antiviral mechanisms of human defensins. J Mol Biol. 2013;425: 4965–80. doi:10.1016/j.jmb.2013.09.038

21. Brice DC, Diamond G. Antiviral Activities of Human Host Defense Peptides. Curr Med Chem. 2020;27: 1420–1443. doi:10.2174/0929867326666190805151654

22. Xu D, Lu W. Defensins: A Double-Edged Sword in Host Immunity. Front Immunol. 2020;11: 764. doi:10.3389/fimmu.2020.00764

23. Zhao C, Porter JM, Burke PC, Arnberg N, Smith JG. Alpha-defensin binding expands human adenovirus tropism. PLoS Pathog. 2024;20: e1012317. doi:10.1371/journal.ppat.1012317

24. Gounder AP, Wiens ME, Wilson SS, Lu W, Smith JG. Critical determinants of human alpha-defensin 5 activity against non-enveloped viruses. J Biol Chem. 2012;287: 24554–62. doi:10.1074/jbc.M112.354068

25. Tenge VR, Gounder AP, Wiens ME, Lu W, Smith JG. Delineation of interfaces on human alpha-defensins critical for human adenovirus and human papillomavirus inhibition. PLoS Pathog. 2014;10: e1004360. doi:10.1371/journal.ppat.1004360

26. Nemerow GR, Stewart PL. Role of alpha(v) integrins in adenovirus cell entry and gene delivery. Microbiol Mol Biol Rev. 1999;63: 725–34.

27. Smith JG, Wiethoff CM, Stewart PL, Nemerow GR. Adenovirus. Curr Top Microbiol Immunol. 2010;343: 195–224. doi:10.1007/82_2010_16

28. Dun Y, Yan J, Wang M, Wang M, Liu L, Yu R, et al. Rac1-dependent endocytosis and Rab5-dependent intracellular trafficking are required by enterovirus A71 and coxsackievirus A10 to establish infections. Biochem Biophys Res Commun. 2020;529: 97–103. doi:10.1016/j.bbrc.2020.05.058

29. Yeh-Kai L, Akusjarvi G, Alestrom P, Pettersson U, Tremblay M, Weber J. Genetic identification of an endoproteinase encoded by the adenovirus genome. J Mol Biol. 1983;167: 217–22.

30. Imelli N, Ruzsics Z, Puntener D, Gastaldelli M, Greber UF. Genetic reconstitution of the human adenovirus type 2 temperature-sensitive 1 mutant defective in endosomal escape. Virol J. 2009;6: 174. doi:1743-422X-6-174 [pii] 10.1186/1743-422X-6-174

31. Perez-Berna AJ, Marabini R, Scheres SH, Menendez-Conejero R, Dmitriev IP, Curiel DT, et al. Structure and uncoating of immature adenovirus. J Mol Biol. 2009;392: 547–57. doi:S0022-2836(09)00787-6 [pii] 10.1016/j.jmb.2009.06.057

32. Silvestry M, Lindert S, Smith JG, Maier O, Wiethoff CM, Nemerow GR, et al. Cryo-electron microscopy structure of adenovirus type 2 temperature-sensitive mutant 1 reveals insight into the cell entry defect. J Virol. 2009;83: 7375–83. doi:10.1128/JVI.00331-09

33. Wiethoff CM, Wodrich H, Gerace L, Nemerow GR. Adenovirus protein VI mediates membrane disruption following capsid disassembly. J Virol. 2005;79: 1992–2000.

34. Scherer J, Yi J, Vallee RB. Role of cytoplasmic dynein and kinesins in adenovirus transport. FEBS Lett. 2020;594: 1838–1847. doi:10.1002/1873-3468.13777

35. Roossien DH, Miller KE, Gallo G. Ciliobrevins as tools for studying dynein motor function. Front Cell Neurosci. 2015;9: 252. doi:10.3389/fncel.2015.00252

36. Liu Q, Long J-E. Insight into the life cycle of enterovirus-A71. Viruses. 2025;17: 181. doi:10.3390/v17020181

37. Strunze S, Trotman LC, Boucke K, Greber UF. Nuclear targeting of adenovirus type 2 requires CRM1-mediated nuclear export. Mol Biol Cell. 2005;16: 2999–3009.

38. Bailey CJ, Crystal RG, Leopold PL. Association of adenovirus with the microtubule organizing center. J Virol. 2003;77: 13275–87.

39. Wang I-H, Burckhardt CJ, Yakimovich A, Morf MK, Greber UF. The nuclear export factor CRM1 controls juxta-nuclear microtubule-dependent virus transport. J Cell Sci. 2017;130: 2185–2195. doi:10.1242/jcs.203794

40. Arias CF, Lopez S. Rotavirus cell entry: not so simple after all. Curr Opin Virol. 2021;48: 42–48. doi:10.1016/j.coviro.2021.03.011

41. de Leeuw E, Li C, Zeng P, Diepeveen-de Buin M, Lu WY, Breukink E, et al. Functional interaction of human neutrophil peptide-1 with the cell wall precursor lipid II. FEBS Lett. 2010;584: 1543–8. doi:S0014-5793(10)00198-5 [pii] 10.1016/j.febslet.2010.03.004

42. Wanniarachchi YA, Kaczmarek P, Wan A, Nolan EM. Human defensin 5 disulfide array mutants: disulfide bond deletion attenuates antibacterial activity against Staphylococcus aureus. Biochemistry. 2011;50: 8005–17. doi:10.1021/bi201043j

43. Chileveru HR, Lim SA, Chairatana P, Wommack AJ, Chiang I-L, Nolan EM. Visualizing attack of escherichia coli by the antimicrobial peptide human defensin 5. Biochemistry. 2015;54: 1767–1777. doi:10.1021/bi501483q

44. Ouellette AJ. Paneth cell alpha-defensins in enteric innate immunity. Cell Mol Life Sci. 2011;68: 2215–29. doi:10.1007/s00018-011-0714-6

45. Hristova K, Selsted ME, White SH. Interactions of monomeric rabbit neutrophil defensins with bilayers: comparison with dimeric human defensin HNP-2. Biochemistry. 1996;35: 11888–94. doi:10.1021/bi961100d

46. Satchell DP, Sheynis T, Shirafuji Y, Kolusheva S, Ouellette AJ, Jelinek R. Interactions of mouse Paneth cell alpha-defensins and alpha-defensin precursors with membranes. Prosegment inhibition of peptide association with biomimetic membranes. J Biol Chem. 2003;278: 13838–46. doi:10.1074/jbc.M212115200 M212115200 [pii]

47. Kagan BL, Selsted ME, Ganz T, Lehrer RI. Antimicrobial defensin peptides form voltage-dependent ion-permeable channels in planar lipid bilayer membranes. Proc Natl Acad Sci U A. 1990;87: 210–4. doi:10.1073/pnas.87.1.210

48. White SH, Wimley WC, Selsted ME. Structure, function, and membrane integration of defensins. Curr Opin Struct Biol. 1995;5: 521–527. doi:10.1016/0959-440X(95)80038-7

49. Maier O, Galan DL, Wodrich H, Wiethoff CM. An N-terminal domain of adenovirus protein VI fragments membranes by inducing positive membrane curvature. Virology. 2010. doi:S0042-6822(10)00226-6 [pii] 10.1016/j.virol.2010.03.043

50. Burckhardt CJ, Suomalainen M, Schoenenberger P, Boucke K, Hemmi S, Greber UF. Drifting motions of the adenovirus receptor CAR and immobile integrins initiate virus uncoating and membrane lytic protein exposure. Cell Host Microbe. 2011;10: 105–17. doi:10.1016/j.chom.2011.07.006

51. Vogt D, Zaver S, Ranjan A, DiMaio T, Gounder AP, Smith JG, et al. STING is dispensable during KSHV infection of primary endothelial cells. Virology. 2020;540: 150–159. doi:10.1016/j.virol.2019.11.012

52. Smith JG, Cassany A, Gerace L, Ralston R, Nemerow GR. Neutralizing antibody blocks adenovirus infection by arresting microtubule-dependent cytoplasmic transport. J Virol. 2008;82: 6492–500. doi:10.1128/JVI.00557-08

53. Pachitariu M, Stringer C. Cellpose 2.0: how to train your own model. Nat Methods. 2022;19: 1634–1641. doi:10.1038/s41592-022-01663-4

54. Stirling DR, Swain-Bowden MJ, Lucas AM, Carpenter AE, Cimini BA, Goodman A. CellProfiler 4: improvements in speed, utility and usability. BMC Bioinformatics. 2021;22: 433. doi:10.1186/s12859-021-04344-9

